# Rapid changes in wing size and shape during the transition to laboratory conditions in a Neotropical drosophilid

**DOI:** 10.64898/2026.09.03.749121

**Authors:** Luísa de Paula Bouzada Dias, Marina Magalhães Moreira, Letícia Carlesso de Paula Sena, Karla Yotoko

## Abstract

The establishment of laboratory populations provides an opportunity to document rapid phenotypic changes across generations, although the processes underlying these changes may be difficult to disentangle. Using geometric morphometrics, we investigated changes in wing size and shape in the Neotropical drosophilid *Drosophila sturtevanti*. We compared field-collected males (Field) with descendants from 12 isofemale lines established independently from field-collected females, in the first (F1) and tenth (F10) laboratory generations. Wing size showed heterogeneous responses in F1, followed by a consistent reduction across all lines by F10, possibly reflecting differences among founders and developmental conditions, including larval density and temperature. Wing shape changed significantly in every line between Field and F1 and between F1 and F10. Changes in wing shape followed broadly similar directions from Field to F1 but became more heterogeneous among lines from F1 to F10. Total shape variance increased significantly between Field and F1. Within-line shape variation was not significantly lower in F1 or F10 than the variation observed in Field, while between-line shape variance increased from F1 to F10. These patterns are consistent with the possible expression of previously cryptic genetic variation following environmental change and the exposure of phenotypic effects of pre-existing genetic variation as homozygosity increases within isolated lines. Our results demonstrate that substantial morphological changes can be detected within the first laboratory generation and continue during subsequent maintenance, highlighting the potential of geometric morphometrics of insect wings for documenting rapid phenotypic change.

## Introduction

Studies of biological invasions, as well as those of populations colonizing novel environments, have provided opportunities to witness “evolution in action”, revealing that marked changes can occur within just a few generations (e.g., Grant & Grant, 1997; Keller & Taylor, 2008; Sherpa et al., 2023). Laboratory studies offer another opportunity to investigate evolutionary dynamics over short timescales, an approach widely used in experimental evolution with microorganisms and viruses (Ascensao & Desai, 2026; Sachdeva et al., 2026).

Among animals, *Drosophila* species are particularly suitable for such investigations because their short generation times make it possible to examine the early stages of laboratory maintenance directly. Studies examining the early generations of laboratory maintenance have documented rapid changes in Behaviour (Stanley & Kulathinal, 2016) and life-history traits, including development time and fecundity (Santos et al., 2012; Simões et al., 2007). A widely used approach in *Drosophila* research is the establishment of isofemale lines, each founded by a single wild-caught, previously inseminated female (David et al., 2005). These lines allow the descendants of different founders to be followed separately, but they also combine an abrupt environmental transition with severe founder sampling and increasing relatedness across generations. In addition, the first laboratory generations may retain maternal effects associated with the environmental conditions experienced by the founding females in the wild (Matos et al., 2002). In the absence of gene flow among lines, genetic drift, inbreeding, and selection may lead to distinct evolutionary trajectories, potentially increasing phenotypic differentiation among lines.

Insect wings provide a particularly suitable system for investigating phenotypic changes during the early stages of laboratory establishment in *Drosophila*. Geometric morphometrics allows size and shape variation to be quantified as separate components while preserving the spatial relationships among homologous landmarks (Bookstein, 1996). Dipteran wings are nearly planar and contain a network of readily identifiable vein intersections, allowing subtle morphological differences to be measured with high precision (Houle et al., 2017; Tan et al., 2026). We applied this approach to *Drosophila sturtevanti* Duda, 1927, a widespread Neotropical drosophilid that can be locally abundant and whose populations harbor substantial genetic variation, much of it distributed within populations rather than among them (Trava et al., 2021; Zorzato et al., 2021).

In this study, we investigated changes in the wing size and shape of male *D. sturtevanti* during the establishment and subsequent maintenance of isofemale lines under laboratory conditions, following our standard laboratory rearing protocol. We established 12 isofemale lines from field-collected females and used geometric morphometrics to quantify variation in wing size and shape. Specifically, we compared a field sample of males (Field) with those from the first laboratory generation (F1) to characterize differences in wing morphology associated with the transition from the field to the laboratory. We then compared F1 and F10 males to examine how these differences developed over subsequent generations, during which maternal effects may have diminished while the effects of genetic drift, inbreeding, and laboratory rearing conditions may have accumulated.

## 1. Materials and Methods

### 1.1. Field sampling and establishment of isofemale lines

In April 2022, banana-baited traps described by Medeiros & Klaczko (1999) were placed in an Atlantic Forest fragment in Minas Gerais, southeastern Brazil (20°45′36.2″ S, 42°52′14.6″ W). The traps remained in the field for 48 h. After collection, specimens were transported to the laboratory, where they were anaesthetized with carbon dioxide and sorted.

Flies that potentially belonged to the *Drosophila saltans* group were initially recognized based on external morphological characteristics, including their generally dark coloration (Yassin et al., 2011). Field-collected males were immediately preserved in absolute ethanol and stored at −20 °C for subsequent species identification. Because reliable identification within the group is based primarily on the morphology of the male aedeagus (Magalhães, 1962), these males were later dissected and identified. Those assigned to *D. sturtevanti* constituted the field sample, hereafter referred to as Field.

Field-collected females potentially belonging to the *D. saltans* group were individually placed in flat-bottomed culture vials (8.1 cm high × 2.5 cm in diameter) containing approximately 2.5 cm of banana–barley medium (Moreira et al., 2026) to establish isofemale lines. Because no males were added to the vials, the offspring originated from matings that occurred before the females were individually isolated in the experimental vials and constituted the first laboratory generation (F1). At least one male from each isofemale line was dissected for aedeagus examination and species identification. Thirteen isofemale lines of *D. sturtevanti* were initially established, of which 12 were maintained until F10 and included in the comparisons between laboratory generations.

### 1.2. Maintenance of isofemale lines

Vials containing field-collected D. sturtevanti females were inspected every two days for the presence of larvae. Once the first larvae were detected in a given vial, the corresponding founding female was preserved in absolute ethanol and stored at −20 °C. As F1 adults emerged, newly emerged individuals were transferred every two days to the same culture vial within each line, where they were allowed to mate freely and reproduce. Once the first F2 larvae were detected in a given line, its F1 adults were preserved in absolute ethanol and stored at −20 °C, and the vials containing F2 larvae were monitored for adult emergence. The same procedure was repeated until F10, with all emerging adults from each generation transferred to a single vial per line. This procedure minimized generation overlap. Developmental times varied among lines, but generations were counted independently for each line, according to its own developmental and reproductive timing. Neither adult nor larval density was standardized at any generation, and eggs and larvae were not transferred between vials.

Following the standard procedure used in our laboratory when incorporating newly established isofemale lines into the stock collection, the lines were maintained at room temperature during the first four generations and kept separately from other laboratory stocks to reduce the risk of introducing ectoparasites, fungi, or other contaminants originating from the field (University of Cambridge Fly Facility, n.d.). From F5 onward, the lines were maintained in a climate-controlled incubator at 23 °C under a 12 h light:12 h dark photoperiod.

### 1.3. Wing preparation and landmark digitization

Morphometric analyses were restricted to the right wings of males from Field, F1, and F10. Nineteen *D. sturtevanti* males were collected in the field, but only 17 were included in the morphometric analyses because the right wings of two individuals were damaged and did not retain all anatomical landmarks. We aimed to analyze 30 males per line in both generations. However, in F1, sample sizes ranged from 13 to 30 because some lines produced fewer than 30 males. In F10, 30 males were analyzed from each of the 12 lines, resulting in a total of 644 individuals across all groups.

Before dissection, ethanol-preserved specimens were rehydrated in 1× phosphate-buffered saline (PBS; pH 7.4). Wing laterality was determined with the fly positioned in anatomical orientation, dorsal side up and with the head facing forward. The wings were then removed at their insertion into the thorax and mounted in their corresponding anatomical positions on a microscope slide. Each image included both wings from a single individual and a 1-mm scale for estimating wing size. Images were obtained using a Nikon SMZ 745T stereomicroscope equipped with a Labomed iVu 3100 digital camera.

Following Prestes et al. (2023), twelve anatomical landmarks were defined at intersections between wing veins or between a vein and the wing margin (Figure 1). Specimens with damaged or deformed wings that prevented the reliable identification of all 12 landmarks were excluded from the analyses.

**Figure 1.**
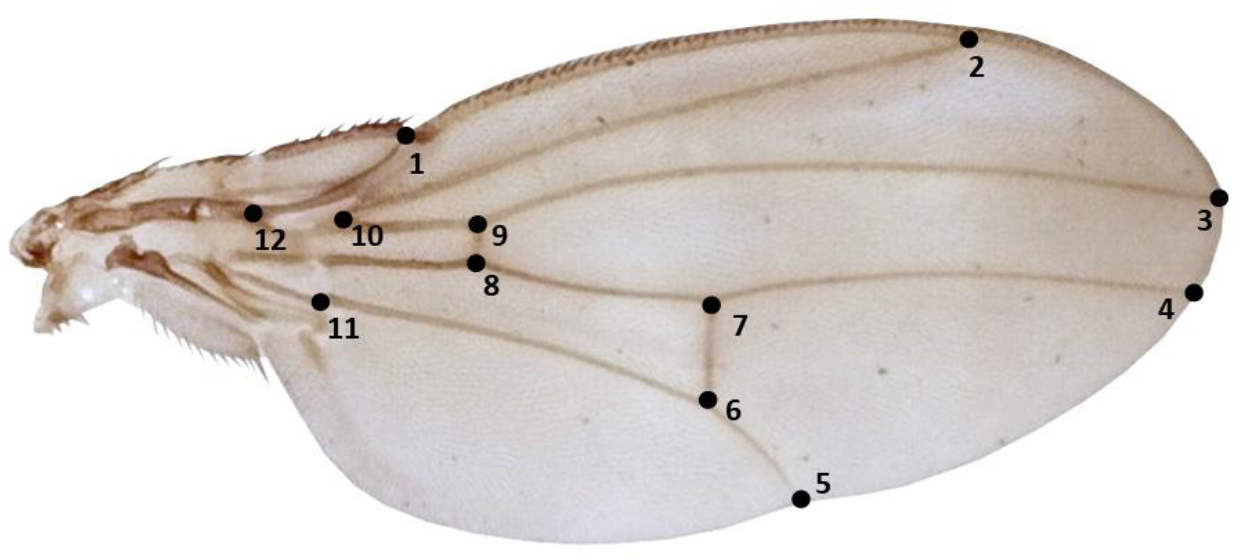
Right wing of a male *Drosophila sturtevanti* in dorsal view, showing the 12 landmarks used in the geometric morphometric analyses.

Landmarks and the scale were manually digitized by a single operator using tpsDig v.2.32 (Rohlf, 2015), which converted their positions into Cartesian coordinates. Each wing was digitized independently twice, always following the same landmark sequence, to quantify digitizing error. The resulting TPS file was subsequently imported into R.

### 1.4. Statistical environment and R packages

All statistical analyses were performed in R (R Core Team, 2026). Geometric morphometric analyses were conducted using the geomorph package (Adams et al., 2026; Baken et al., 2021), with residual-randomization procedures implemented using RRPP (Collyer & Adams, 2018). Estimated marginal means and planned contrasts were obtained using emmeans (Lenth & Piaskowski, 2017), Brown– Forsythe tests were implemented using the car package (Fox J & Weisberg S, 2019), and data manipulation and visualization were conducted using dplyr and ggplot2 (Wickham, 2016; Wickham et al., 2026).

### 1.5. Assessment of digitizing error

Digitizing errors were evaluated separately for wing shape and centroid size. To assess digitizing error in wing shape, all landmark configurations from both digitizing replicates were subjected to a Generalized Procrustes Analysis (GPA) using the gpagen function of the geomorph package. This digitizing-error GPA was used only to assess digitizing error, which was evaluated using a Procrustes linear model fitted with the procD.lm function:

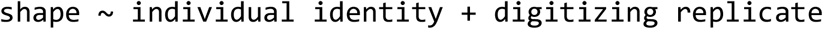

Individual identity represented biological variation among wings, whereas digitizing replicate tested whether the two landmarking rounds differed systematically. Residual variation represented random differences between repeated digitizations of the same wing that were not attributable to a systematic replicate effect.

To assess digitizing error in centroid size, a conventional linear model was fitted to log-transformed centroid size, with individual identity and digitizing replicate as explanatory variables:

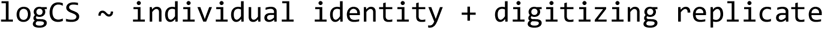

After digitizing error was assessed, the Procrustes coordinates obtained for the two digitizing replicates of each wing were averaged landmark by landmark, producing a single mean Procrustes configuration per individual. These mean configurations were then jointly subjected to a second Generalized Procrustes Analysis, hereafter referred to as the biological-analysis GPA. The Procrustes coordinates resulting from this second GPA were used in all subsequent analyses of wing shape. For centroid size, the two values obtained from the digitizing-error GPA were averaged for each individual, and the natural logarithm of this mean centroid size was used in the statistical analyses.

### 1.6. Framework for pairwise comparisons

The analyses included 25 biological groups: one field group (Field), 12 F1 groups, one for each isofemale line, and 12 corresponding F10 groups. Pairwise comparisons of mean centroid size and mean wing shape followed the same planned framework. To investigate whether the transition from field to laboratory conditions was associated with changes in wing size and shape already in the first generation, each F1 line was compared separately with Field. This approach allowed us to evaluate the initial phenotypic differentiation of each independently established line from the field reference. To evaluate subsequent changes during laboratory maintenance, F1 and F10 were compared within each line.

The planned comparisons comprised two sets of 12 tests: Field – F1 and F1 – F10. The two sets were treated as separate families of tests, and P-values were adjusted using the Benjamini–Hochberg procedure (Benjamini & Hochberg, 1995) implemented with the p.adjust function in R (method = “BH”).

### 1.7. Centroid-size analyses

Centroid size, defined as the square root of the summed squared distances of all landmarks from their centroid, was used as the measure of wing size (Zelditch et al., 2012).

Differences in mean centroid size were evaluated using linear models (section 2.6). For the Field–F1 analyses, the model included a categorical factor distinguishing the field sample from F1 of each isofemale line (log centroid size ∼ group). For the F1–F10 analyses, the model included generation, line, and their interaction, allowing us to test whether the magnitude of change between generations differed among isofemale lines:

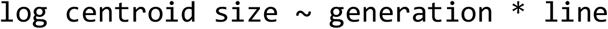

In both analyses, planned comparisons were obtained from estimated marginal means using the emmeans package.

In addition to differences in mean size, differences in the variance of centroid size were assessed using Brown–Forsythe tests, implemented as median-centered Levene tests with the leveneTest function of the car package.

### 1.8. Wing-shape analyses

#### 1.8.1. Tests of Allometry

To assess the role of allometry in wing-shape variation, we performed two complementary Procrustes ANOVAs. First, we analyzed all 644 individuals, treating Field and each isofemale line in F1 and F10 as 25 distinct biological groups. The model included log-transformed centroid size, group, and their interaction to test whether allometric relationships differed among groups. Second, we analyzed only F1 and F10, treating generation and line as crossed factors because the same 12 lines were represented in both generations. This model included log-transformed centroid size, generation, line, and all two-way and three-way interactions, allowing us to test whether allometric relationships differed between generations, among lines, and in their changes across generations.

#### 1.8.2. Principal Component Analyses

The Procrustes coordinates obtained from the biological-analysis GPA (section 2.5) were used as shape variables in the statistical analyses and in a principal component analysis conducted to visualize the distribution of observed wing-shape variation among Field and the F1 and F10 groups. The allometric analysis described in section 2.8.1 was used to determine whether a single allometric correction would be appropriate. If allometric trajectories did not differ among generations or among lines within generations, a global allometric correction would be applied before the PCA. Otherwise, the PCA would be performed on the original Procrustes coordinates, without removing the allometric component, to avoid introducing distortions in groups with different size–shape relationships (Klingenberg, 2016). Thus, when no common allometric trajectory was supported, the PCA represented the total observed variation in wing shape, including its allometric component, defined as shape variation associated with size (Klingenberg, 2016).

#### 1.8.3. Planned pairwise comparisons of mean shape

Differences in mean wing shape were evaluated separately for each planned comparison (section 2.6) using Procrustes linear models fitted with the procD.lm function. These pairwise analyses accounted for allometry by comparing the predicted mean shapes of each pair of groups at a common reference size. For each compared pair, log-transformed centroid size was included as a covariate, together with its interaction with group. The interaction tested whether the two groups differed in their allometric trajectories.

Before fitting each model, we examined the observed range of log-transformed centroid size in the two groups to select an appropriate reference size. When the joint mean size fell within the observed range of both groups, it was used as the reference size. When the joint mean fell outside the range of one group, but the size ranges overlapped, the midpoint of the shared range was used instead. When the size ranges did not overlap, the joint mean was retained as the reference size, and the resulting comparison was considered extrapolative.

For each comparison, log-transformed centroid size was centered on the selected reference size, and the following model was fitted:

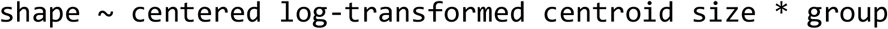

The group effect tested the difference between the shapes predicted for the two groups at the selected common reference size, whereas the interaction tested whether their allometric trajectories differed.

For each comparison, the Procrustes distance between the two shapes predicted at the common reference size was calculated as a measure of the magnitude of shape differentiation.

#### 1.8.4. Analyses of shape variance

Total wing-shape variation was quantified as Procrustes variance calculated from the coordinates obtained from the biological-analysis GPA. Because these coordinates describe shape after the removal of differences in position, orientation, and scale, Procrustes variance represents variation in wing shape rather than variation in centroid size. The allometric analysis described in section 2.8.1 was used to determine whether a common allometric correction would be appropriate. If a common allometric trajectory was supported, Procrustes variance would be calculated from allometry-corrected shape coordinates. Otherwise, variance would be calculated from the original Procrustes coordinates, thereby representing total observed shape variation, including the component of shape variation associated with centroid size.

Total Procrustes variance was calculated separately for Field, pooled F1, and pooled F10, using all available individuals. To assess uncertainty and facilitate comparisons using standardized sample sizes, we additionally performed bootstrap resampling. For each dataset, 17 individuals were sampled with replacement in each of 9,999 bootstrap replicates, corresponding to the number of field-collected males. Pairwise comparisons among Field, pooled F1, and pooled F10 were based on the resulting bootstrap distributions of variance differences and ratios.

For F1 and F10, the observed total Procrustes variance was further partitioned into within-line and between-line components. The within-line component was calculated from the squared Procrustes deviations of individuals from the mean shape of their respective line. The between-line component was calculated from the squared Procrustes distances between each line mean and the overall mean shape of the generation, weighted by the number of individuals in each line. Both components were expressed using the same denominator as total Procrustes variance, so that their sum equaled the total variance. This decomposition was used to determine whether changes in total shape variation arose primarily from variation among individuals within lines or from divergence among isofemale lines.

To assess whether differences in sample sizes between F1 and F10 influenced the observed variance components, we performed a sensitivity analysis using 9,999 subsampling replicates. In each replicate, 13 individuals were randomly selected without replacement from each of the 12 isofemale lines in both generations, corresponding to the smallest line-specific sample size in F1 (N = 156 individuals per generation). Within-line and between-line variance components were recalculated using the same procedure applied to the complete dataset, and the distribution of between-line variance ratios (F10/F1) was examined.

## 2. Results

### 2.1. Assessment of digitizing error

Digitizing replicate had a statistically significant effect on both centroid size (*F*_1,643_ = 21.16, *P* = 5.10 × 10^−6^) and wing shape (*F*_1,643_ = 63.77, *Z* = 8.48, *P* = 0.001) (Tables S1 and S2). However, it accounted for only 0.04% of the total variation in centroid size and 0.82% of the total variation in wing shape. In contrast, differences among individuals accounted for approximately 98.72% and 90.89% of the variation in centroid size and shape, respectively, while residual variation accounted for 1.24% and 8.29%. Thus, although statistically significant, the effect of digitizing replicate was small relative to the among-individual variation of primary biological interest in this study, supporting the use of the landmark data in subsequent analyses (Cardini, 2025). To further reduce their influence, the two digitizations were averaged landmark by landmark for each individual before the subsequent analyses, as described in the Methods.

### 2.2. Centroid-size analyses

When right-wing centroid size was compared between the field sample (Field) and F1, seven of the 12 isofemale lines exhibited significantly different wing sizes relative to the field sample: six lines had smaller wings than Field (L01, L06, L13, L14, L22, and L23), whereas L15 had larger wings (Figure 2). No significant differences were detected for the remaining five lines. In the F1–F10 comparisons, centroid size decreased significantly in all 12 lines (BH-adjusted *P* < 0.05; Figure 2). Across the 12 isofemale lines, mean wing centroid size decreased by 9.2% from F1 to F10, with reductions ranging from 3.1% to 17.5% among lines. Full results of the planned contrasts are provided in Supplementary Tables S3 and S4.

**Figure 2.**
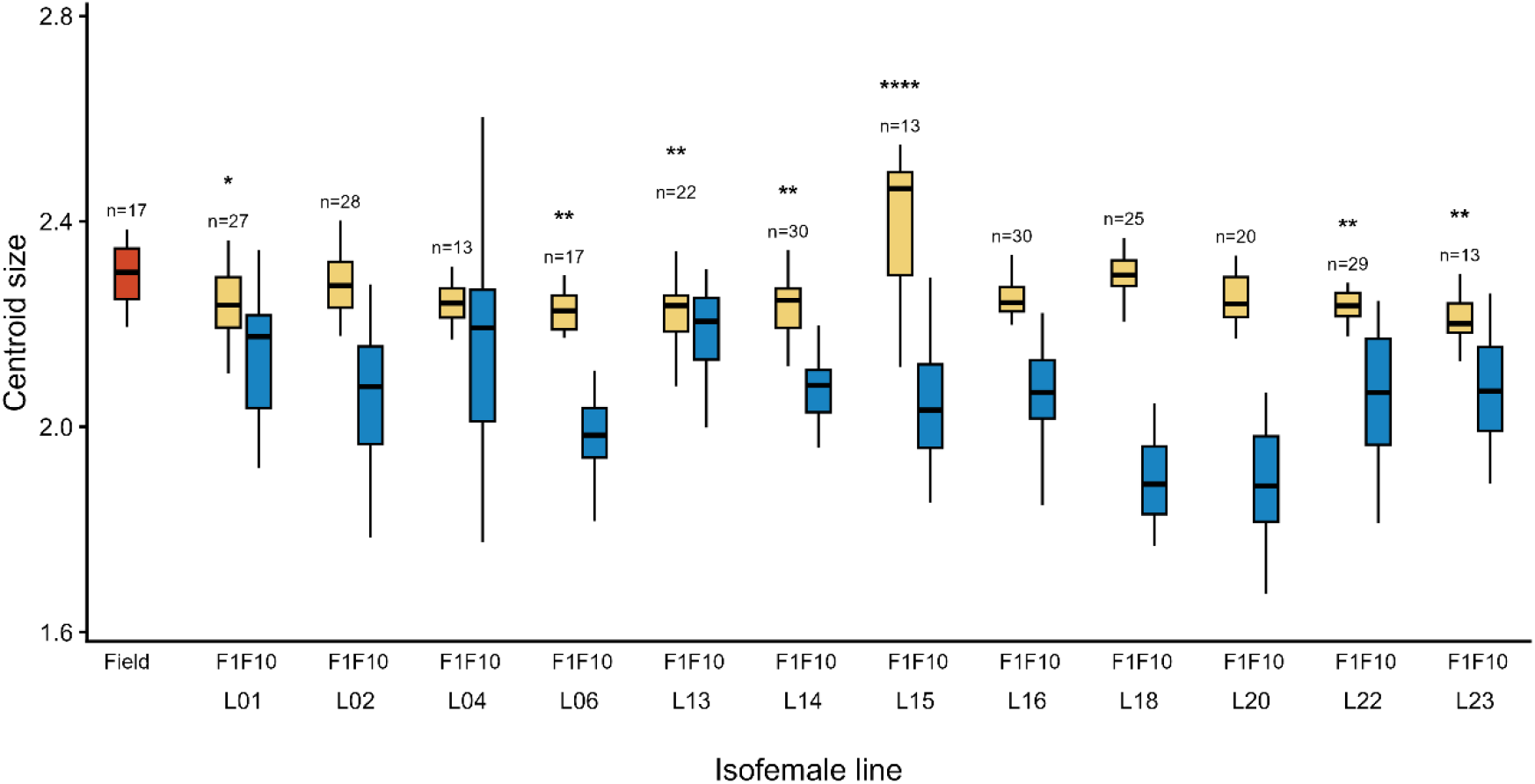
Boxplots of untransformed centroid size in field-collected males (Field, red) and in males from the first (F1, yellow) and tenth (F10, blue) laboratory generations of 12 isofemale lines of *Drosophila sturtevanti*. Asterisks above the F1 boxplots indicate significant differences between F1 and Field after Benjamini–Hochberg correction (Table S3). Centroid size was significantly smaller in F10 than in F1 in all isofemale lines (maximum Benjamini–Hochberg-adjusted *P* = 0.0156; Table S4). Numbers above the Field and F1 boxplots indicate sample sizes; all F10 groups comprised 30 individuals. Statistical analyses were performed using log-transformed centroid size. Significance levels are indicated as follows: *P* < 0.05 (\**), P* < 0.01 *(**),* and *P* < 0.0001 (****).

No significant differences in the variance of log-transformed centroid size were detected between Field and F1 for any isofemale line after Benjamini– Hochberg correction (Table S5). In contrast, variance increased significantly from F1 to F10 in eight of the 12 isofemale lines after correction (L02, L04, L06, L16, L18, L20, L22, and L23; Table S6). In three of the four remaining lines, variance in F10 was still more than twice that observed in F1, whereas L15 showed little change

### 2.3. Wing-shape analyses

The Procrustes ANOVA including all 25 biological groups revealed significant allometry in wing shape, with log-transformed centroid size explaining 7.86% of total shape variation (*F* = 89.88, *Z* = 9.73, *p* = 0.0001). Wing shape differed significantly among groups (*R*^2^ = 0.368, *F* = 17.52, *Z* = 29.24, *p* = 0.0001), and the significant interaction between centroid size and group (*R*^2^ = 0.034, *F* = 1.62, *Z* = 4.89, *p* = 0.0001) indicated heterogeneity in allometric trajectories. The analysis restricted to F1 and F10, with generation and line treated as crossed factors, also revealed significant differences in allometric relationships between generations (*R*^2^ = 0.0024, *F* = 2.72, *p* = 0.0101) and among lines (*R*^2^ = 0.046, *F* = 4.66, *p* = 0.0001). Furthermore, the significant three-way interaction between centroid size, generation, and line (*R*^2^ = 0.016, *F* = 1.62, *p* = 0.0004) indicated that changes in allometric trajectories between F1 and F10 differed among lines.

Given the significant heterogeneity in allometric trajectories, the PCA was performed on the original Procrustes coordinates, without applying a global allometric correction, thereby retaining the total observed wing-shape variation, including its allometric component. Figure 3 shows that the F1 groups from all 12 isofemale lines were displaced relative to Field. Consistent with this pattern, the planned comparisons revealed significant differences in wing shape between Field and each F1 line (*P*_BH_ ≤ 0.0014; Table S7).

**Figure 3.**
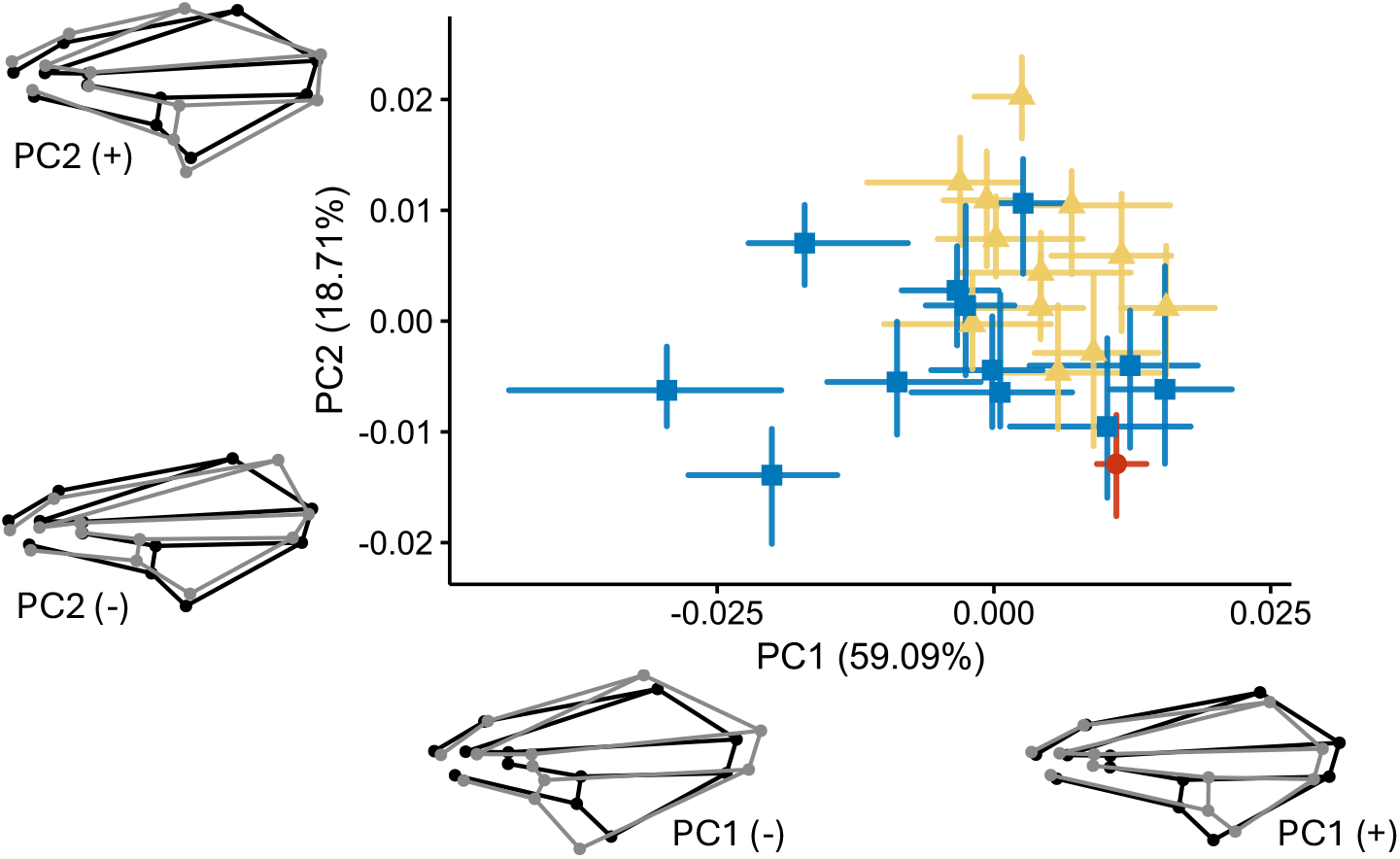
Principal component analysis of wing shape in field-collected males (Field, red circle) and males from the first (F1, yellow triangles) and tenth (F10, blue squares) laboratory generations of 12 isofemale lines of *Drosophila sturtevanti*. Symbols indicate group medians, whereas the horizontal and vertical bars represent the interquartile ranges of PC1 and PC2, respectively. PC1 and PC2 accounted for 59.09% and 18.71% of the total observed variation in wing shape, respectively. Wing shape configurations at the negative and positive extremes of PC1 and PC2 are shown around the plot, with the mean configuration in black and the reconstructed configurations in gray. Shape differences were magnified fourfold to facilitate visualization. Landmark numbering is shown in Figure 1.

Additional changes in wing shape were observed between F1 and F10, with significant differences in all lines (*P*_BH_ ≤ 0.0075; Figure 3; Table S8). However, the centroid-size ranges of L18 and L20 did not overlap between F1 and F10, so the size-adjusted shape comparisons for these lines involved extrapolation. In addition, the allometric trajectories differed significantly between F1 and F10 in L01, L02, L16, and L18.

Beyond the shifts in mean wing shape, we examined whether the transition to the laboratory also changed the amount of shape variation. Because allometric trajectories differed among generations and among lines within generations, Procrustes variance was calculated from the original Procrustes coordinates, without removing a common allometric component, and therefore represents total observed wing-shape variation. When all individuals were considered, Procrustes variance was 39% higher in F1 than in Field, increasing from 4.14 × 10⁻⁴ to 5.77 × 10⁻⁴ (Table S9). The bootstrap analysis confirmed this pattern (Figure 4A), with F1 variance being typically 46% higher than Field variance in samples standardized to 17 individuals per generation (Table S10). Thus, the greater wing-shape variance observed in F1 persisted after standardizing sample sizes and was already detectable in the first laboratory generation.

**Figure 4.**
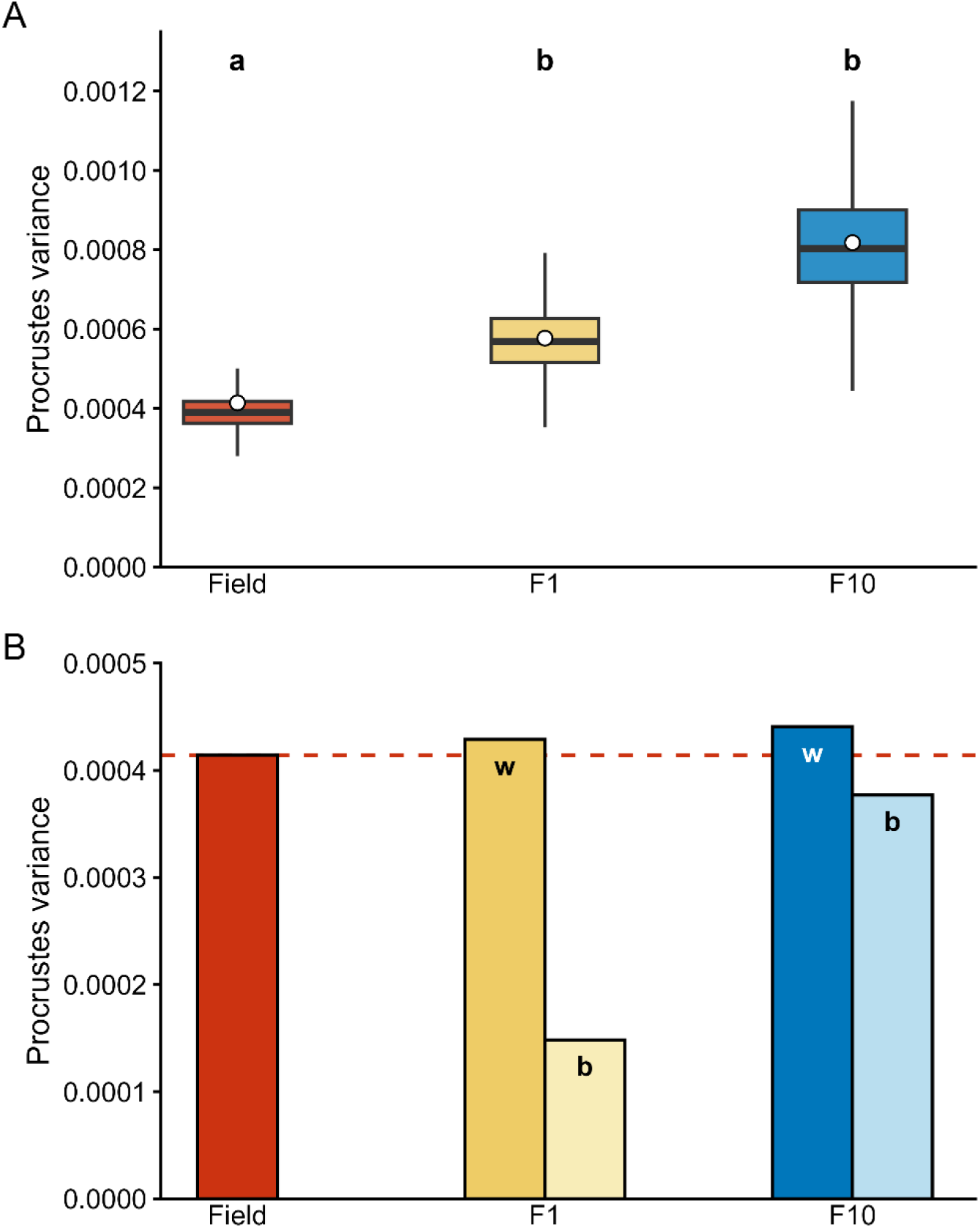
Variation in wing shape across field and laboratory generations of *Drosophila sturtevanti*. (A) Distributions of total Procrustes variance obtained from 9,999 bootstrap samples of 17 individuals drawn with replacement from the field sample (Field) and from the first (F1) and tenth (F10) laboratory generations. White circles indicate the observed Procrustes variance calculated using all individuals available in each dataset. Different lowercase letters above the boxplots indicate differences between datasets based on the 95% bootstrap confidence intervals of pairwise differences (Table S10); groups sharing the same letter did not differ. (B) Decomposition of the observed total Procrustes variance in F1 and F10 into within-line (w) and between-line (b) components. The Field bar represents the total observed variance in the field sample. The dashed red line indicates the observed total variance in Field and is shown as a reference for comparison with the within– and between-line components of the laboratory generations. Within– and between-line components sum to the corresponding total observed variance shown in panel A.

The increase in total variance from Field to F1 did not result simply from the partitioning of field variation among the newly established isofemale lines. Within each line, F1 variance did not differ significantly from that observed in Field, despite each F1 sample consisting of descendants of a single founding female (Table S11). At the same time, differentiation among lines added a between-line component to total variance in F1 (Figure 4B; Table S12). Thus, the increase in total variance in the first laboratory generation reflected both the absence of a detectable reduction in variation among descendants within lines and the emergence of variation among lines.

Within-line variance also did not differ significantly between F1 and F10 in any of the 12 lines, providing no evidence of a decline in phenotypic shape variance during the subsequent generations of laboratory maintenance (Table S12). In contrast, the between-line component was larger in F10 than in F1 (Figure 4B; Table S12). A sensitivity analysis showed that this pattern persisted when sample sizes were standardized across lines and generations, indicating that the increase in between-line variance was not simply a consequence of the smaller sample sizes of some lines in F1 (Table S13).

## 3. Discussion

We investigated changes in wing size and shape during the establishment and subsequent maintenance of *Drosophila sturtevanti* isofemale lines under routine laboratory conditions. Our results show that both wing size and shape changed rapidly, although the patterns differed between traits and across generations. Wing size showed heterogeneous changes among lines from Field to F1, followed by a consistent reduction from F1 to F10. In contrast, wing shape changed in broadly similar directions across the 12 lines during the transition from Field to F1, with all lines exhibiting significant differences from the Field sample. From F1 to F10, wing-shape changes became more divergent among lines, resulting in increased between-line differentiation.

The heterogeneous responses in wing size observed during the first stage (Field – F1) may reflect differences among the founding females and the genetic backgrounds represented by each line. Because each line originated from a single field-collected female, variation among lines may have arisen from maternal effects (Wolf & Wade, 2009), including differences in founder age and associated effects on egg provisioning (Kern et al., 2001). Additional sources of variation include the sampling of distinct maternal and paternal genetic backgrounds from the natural population, with possible variation in the number and identity of males that had mated with each founding female before capture (David et al., 2005; Hoffmann & Parsons, 1998), and genotype-specific reaction norms expressed in the novel laboratory environment (Via & Lande, 1985).

Although larval density was not explicitly controlled, it was probably lower during the production of F1 than during the production of F10. F1 was produced by a single founder female, whereas F10 was produced by multiple females reproducing within each line, potentially resulting in greater egg production in F10. Consistent with this difference, some F1 lines produced fewer than the target of 30 males, resulting in smaller sample sizes for those lines. Competition for nutrients and oxygen may therefore have been more intense in F10, providing a plausible explanation for the consistent reduction in wing size across lines (Nicholls & Shingleton, 2026). Importantly, this reduction was accompanied by greater size variation in most lines. Rather than affecting all larvae uniformly, crowding may have generated unequal access to resources, allowing some individuals to grow relatively well while others experienced stronger nutritional limitation and developed substantially smaller wings (Bakker, 1962).

Developmental temperature may also have contributed to the larger wing size observed in F1 than in F10, because the first laboratory generations were maintained outside the temperature-controlled chamber during a relatively cool period and the temperatures experienced by the flies were not recorded (Thomas, 1993). The lack of temperature control during the first laboratory generations limits our ability to identify the factors responsible for the observed changes in wing size. Nevertheless, these conditions reflect the routine procedures used in our laboratory to establish new isofemale lines.

Wing shape also followed a stage-dependent pattern. All isofemale lines exhibited significant differences in wing shape in F1 relative to the Field sample, and total shape variance across the 12 lines was 39% higher in F1 than in Field. This increase did not merely reflect divergence among the newly established lines. Despite each line comprising descendants of a single field-collected female, within-line variance was not detectably lower than the variance observed in Field in any of the 12 lines. Differentiation among lines therefore added a between-line component without a corresponding detectable contraction of variation among descendants within lines. Thus, the descendants of the field-collected females differed from the Field sample in mean wing shape and occupied a broader region of morphospace.

Together, the rapid displacement of mean shape and the persistence of substantial within-line variation are consistent with an episode of developmental decanalization. Changes in developmental temperature may have contributed to this pattern. Developmental temperature is known to alter mean wing shape (Debat et al., 2008) and may also affect the amount and structure of within-group shape variation (Debat et al., 2009). As originally proposed by Waddington (1942), decanalization may expose cryptic genetic variation that remains phenotypically silent under the environmental conditions to which a population is adapted but becomes expressed when those conditions change (Tawfeeq et al., 2024).

Wing development may be particularly susceptible to environmental perturbations. In *Drosophila melanogaster*, a large-scale functional screen of 10,920 protein-coding genes showed that nearly 7,000 influenced the development of the wing imaginal disc, often through individually small effects (López-Varea et al., 2021; Ostalé et al., 2023). This extensive genetic complexity may help explain both the sensitivity of wing development to environmental change and the line-specific responses observed here. Because each isofemale line was founded by a different wild-caught female, each presumably captured a distinct subset of the genetic variation present in the source population. Interactions between these genetic backgrounds and the novel laboratory environment may therefore have contributed to the line-specific morphological responses observed here. Under this interpretation, variation in wing shape would emerge from the combined effects of many loci rather than from a small number of variants with large effects (Barton, 2022; Tautz et al., 2026).

The trajectory from F1 to F10 differed from the initial response to laboratory transfer. Although the F1 lines occupied a broader region of morphospace than the Field sample, their distribution appeared relatively restricted compared with F10. Wing shape continued to change significantly in every isofemale line. The PCA showed that, relative to Field, all F1 lines were displaced toward more positive PC2 values, while most also occupied more negative positions along PC1. In contrast, changes from F1 to F10 appeared more heterogeneous, with the lines following different directions in morphospace, particularly along PC1. Total shape variance was also numerically higher in F10 than in F1, but, unlike the increase observed from Field to F1, this further increase was not statistically supported.

The decomposition of shape variance provided a more detailed view of the differences between F1 and F10. The between-line component was larger in F10 than in F1, consistent with the line-specific changes in mean wing shape observed across generations. Progressive inbreeding within each isofemale line is expected to increase homozygosity, potentially exposing phenotypic effects of recessive alleles previously masked in heterozygotes (Robertson, 1951). Changes in allele frequencies and homozygosity may also alter multilocus genetic backgrounds and, through epistatic interactions, modify the phenotypic effects of segregating alleles (Goodnight, 1988; Mackay & Anholt, 2024). These processes could therefore have contributed to the increasing differentiation among lines.

The interpretation of our results should consider the functional importance of wings under natural conditions. Wing morphology is likely constrained by natural selection because of its importance for flight, predator avoidance, and reproductive success. As noted above, the phenotypic shape variance observed in Field was no greater than that within individual isofemale lines, suggesting that the range of wing shapes observed in nature may be more restricted than the range of phenotypes that can be produced during development. Supporting this view, Houle & Fierst (2013) found mutational variation across nearly the entire dimensionality of *Drosophila* wing form, while Houle et al. (2017) showed that this substantial mutational potential contrasts with remarkably slow long-term divergence. More recently, Brändén & De Lisle (2026) provided direct evidence of pervasive stabilizing selection on multivariate wing shape. Together, these findings suggest that stabilizing selection and other constraints may restrict the range of wing phenotypes expressed in natural populations.

Although the transition to laboratory conditions may initially be challenging for field-collected organisms, these conditions may nevertheless relax some of the selective pressures experienced in natural environments (Huey & Rosenzweig, 2009). One example is the occurrence of individuals with conspicuous wing-shape deformities in laboratory lines (e.g., D’Ávila et al., 2008), showing that such phenotypes can occur and remain observable under laboratory conditions. In parallel, developmental decanalization may have increased the expression of previously cryptic phenotypic variation (Flatt, 2005; Paaby & Rockman, 2014). These are distinct processes, but they may act synergistically in laboratory populations: decanalization broadens the range of phenotypes produced, whereas relaxed selection may allow a larger fraction of those phenotypes to survive and persist.

Our study showed that the routine establishment and maintenance of *D. sturtevanti* isofemale lines were accompanied by substantial changes in wing morphology, detectable as early as the first laboratory generation and continuing through F10. These changes may reflect the combined effects of an abrupt environmental transition, founder sampling, and subsequent genetic changes within isolated lines. Environmental changes may expose previously cryptic genetic variation, while increasing homozygosity associated with inbreeding may reveal phenotypic effects of genetic variation already present in the founders. Although the relative contributions of these processes remain unknown, the patterns observed here resemble, in some respects, the rapid phenotypic changes documented during biological invasions and the colonization of new environments. Thus, the establishment and maintenance of isofemale lines, combined with geometric morphometric analyses of insect wings, may provide a useful system for visualizing rapid phenotypic changes across generations, offering a window into what has been termed “evolution in action”, and for investigating their underlying mechanisms.

## 4. Acknowledgements

We thank Dr. Hermes Fonsêca de Medeiros for his valuable suggestions on the statistical analyses. We thank Fernanda Mayrink Campos Pereira for her essential assistance during the collection of drosophilids and line maintenance. LPBD, MMM, and LCPS thank the Coordenação de Aperfeiçoamento de Pessoal de Nível Superior (CAPES) – Finance Code 001 for granting their scholarships. LPBD gratefully acknowledges the financial support of the Conselho Nacional de Desenvolvimento Científico e Tecnológico (CNPq) for granting her master’s scholarship during which this study was conducted. LCPS is grateful to the Fundação de Amparo à Pesquisa do Estado de São Paulo (FAPESP) for her current Ph.D scholarship (FAPESP 2024/23360-7). We also acknowledge the Departamento de Biologia Geral for providing the infrastructure to conduct this study.

## 6. Declaration of AI use

During the preparation of this manuscript, the authors used ChatGPT (OpenAI) interactively to assist with language editing and readability, to propose and troubleshoot R code for statistical analyses and figure preparation, and to help organize analysis scripts and associated README documentation for submission. Code generated with AI assistance was iteratively tested, reviewed and, where appropriate, modified by the authors before use. The authors executed all analyses, examined and verified the resulting outputs, evaluated the appropriateness of the analytical approaches and retained responsibility for the interpretation of the results and all scientific conclusions.

NotebookLM (Google), Consensus and Google Scholar Labs were used to assist with literature discovery. All publications identified using these tools were independently accessed and evaluated by the authors. Grammarly was used for language checking, and Turnitin was used for similarity and plagiarism screening. The authors take full responsibility for the accuracy, originality and scientific content of the manuscript.

