## Supplemental Material for "Rapid changes in wing size and shape during the transition to laboratory conditions in a Neotropical drosophilid"

### Assessment of digitizing error

**Table S1.** Analysis of variance for digitizing error in centroid size, based on a linear model with log-transformed centroid size as the response variable and individual identity and digitizing replicate as explanatory factors. Individual identity represents biological variation among wings, digitizing replicate tests for systematic differences between the two digitizing rounds, and the residual term represents random digitizing error within individuals. Degrees of freedom, sums of squares, mean squares, $R^2$, F-values, and P-values are shown.

| **Effect** | **Df** | **SS** | **MS** | **$R^2$** | **F** | **P** |
|---|---|---|---|---|---|---|
| Individual_ID | 643 | 7.40515 | 0.01152 | 0.98722 | 79.79883 | **2.20E-16** |
| replicate | 1 | 0.00305 | 0.00305 | 0.00040 | 21.15748 | **5.10E-06** |
| Residuals | 643 | 0.09280 | 0.00014 | 0.01237 | | |

**Table S2.** Procrustes ANOVA for digitizing error in wing shape, based on a model with the Procrustes coordinates obtained from the digitizing-error GPA as response variables and individual identity and digitizing replicate as explanatory factors. Individual identity represents biological variation among wings, digitizing replicate tests for systematic differences between the two digitizing rounds, and the residual term represents random digitizing error within individuals. Degrees of freedom, sums of squares, mean squares, $R^2$, F-values, Z-scores, and P-values are shown.

| **Effect** | **Df** | **SS** | **MS** | **$R^2$** | **F** | **Z** | **P** |
|---|---|---|---|---|---|---|---|
| Individual_ID | 643 | 0.95579 | 0.00149 | 0.90893 | 10.97079 | 46.64093 | **1.00E-04** |
| replicate | 1 | 0.00864 | 0.00864 | 0.00822 | 63.77172 | 9.15684 | **1.00E-04** |
| Residuals | 643 | 0.08712 | 0.00014 | 0.08285 | | | |
| Total | 1287 | 1.05155 | | | | | |

### Centroid-size analyses

**Table S3.** Planned contrasts in log-transformed centroid size between the field sample (Field) and each isofemale line in F1, calculated as F1 − Field. Negative estimates indicate smaller wings in F1, whereas positive estimates indicate larger wings in F1. The table shows the estimated contrast, standard error (SE), t-ratio, unadjusted P-value, and Benjamini–Hochberg-adjusted P-value. All contrasts were evaluated with 271 residual degrees of freedom.

| Line | estimate | SE | t.ratio | p.value | p_BH |
|---|---|---|---|---|---|
| L01 | -0.01970 | 8.35E-03 | -2.3626 | 0.01890 | **0.03230** |
| L02 | -0.00470 | 8.29E-03 | -0.5654 | 0.57230 | 0.61480 |
| L04 | -0.02000 | 9.94E-03 | -2.0088 | 0.04550 | 0.06070 |
| L06 | -0.02800 | 9.25E-03 | -3.031 | 0.00270 | **0.00800** |
| L13 | -0.02760 | 8.71E-03 | -3.1692 | 0.00170 | **0.00680** |
| L14 | -0.02360 | 8.19E-03 | -2.8764 | 0.00430 | **0.00870** |
| L15 | 0.04740 | 9.94E-03 | 4.7733 | 2.97E-06 | **3.56E-05** |
| L16 | -0.01680 | 8.19E-03 | -2.0567 | 0.04070 | 0.06070 |
| L18 | 0.00430 | 8.48E-03 | 0.5038 | 0.61480 | 0.61480 |
| L20 | -0.01630 | 8.90E-03 | -1.8363 | 0.06740 | 0.08090 |
| L22 | -0.02380 | 8.24E-03 | -2.891 | 0.00420 | **0.00870** |
| L23 | -0.03770 | 9.94E-03 | -3.7886 | 1.87E-04 | **1.12E-03** |

**Table S4.** Planned contrasts in log-transformed centroid size between the first (F1) and the tenth (F10) generations of each isofemale line, calculated as F1 – F10. All estimates were positive, indicating larger wings in F1. The table shows the estimated contrast, standard error (SE), *t*-ratio, unadjusted *P*-value, and Benjamini–Hochberg-adjusted P-value. All contrasts were evaluated with 603 residual degrees of freedom.

| Line | estimate | SE | t.ratio | p.value | p_BH |
|---|---|---|---|---|---|
| L01 | 0.0504 | 0.0129 | 3.9044 | 1.05E-04 | **1.35E-04** |
| L02 | 0.1029 | 0.0128 | 8.0511 | 4.39E-15 | **1.05E-14** |
| L04 | 0.0425 | 0.0161 | 2.6335 | 8.67E-03 | **9.46E-03** |
| L06 | 0.1217 | 0.0148 | 8.2459 | 1.03E-15 | **3.09E-15** |
| L13 | 0.0331 | 0.0136 | 2.425 | 1.56E-02 | **1.56E-02** |
| L14 | 0.0792 | 0.0126 | 6.3129 | 5.31E-10 | **7.97E-10** |
| L15 | 0.1568 | 0.0161 | 9.7132 | 8.08E-21 | **3.23E-20** |
| L16 | 0.0928 | 0.0126 | 7.3952 | 4.75E-13 | **9.50E-13** |
| L18 | 0.1925 | 0.0132 | 14.6239 | 1.13E-41 | **1.36E-40** |
| L20 | 0.1739 | 0.014 | 12.3902 | 1.44E-31 | **8.62E-31** |
| L22 | 0.0838 | 0.0127 | 6.6198 | 7.96E-11 | **1.36E-10** |
| L23 | 0.0628 | 0.0161 | 3.8883 | 1.12E-04 | **1.35E-04** |

**Table S5.** Brown–Forsythe tests comparing the variance of log-transformed centroid size between the field sample (Field) and F1 of each isofemale line. The variance in Field was 0.001129 and was used as the reference in all comparisons. The table shows the variance in F1, the F1/Field variance ratio, the F-statistic, the unadjusted P-value, and the Benjamini–Hochberg-adjusted P-value. None of the comparisons was significant after correction for multiple testing.

| Line | $s^2$ F1 | F1/Field | F_value | p_value | p_BH |
|---|---|---|---|---|---|
| L01 | 0.00111 | 0.98388 | 0.10086 | 0.75237 | 0.75237 |
| L02 | 0.00073 | 0.6462 | 0.39197 | 0.53457 | 0.64149 |
| L04 | 0.00042 | 0.37357 | 1.79245 | 0.1914 | 0.34783 |
| L06 | 0.00028 | 0.24948 | 3.97209 | 0.05485 | 0.16454 |
| L13 | 0.00103 | 0.91123 | 0.12262 | 0.72819 | 0.75237 |
| L14 | 0.00063 | 0.55812 | 0.83082 | 0.36689 | 0.48919 |
| L15 | 0.00338 | 2.99071 | 2.17463 | 0.15146 | 0.34783 |
| L16 | 0.0004 | 0.35218 | 4.23011 | 0.04554 | 0.16454 |
| L18 | 0.0003 | 0.2641 | 5.73113 | 0.02145 | 0.12867 |
| L20 | 0.00047 | 0.41806 | 1.68385 | 0.2029 | 0.34783 |
| L22 | 0.00024 | 0.20906 | 8.66091 | 0.00517 | 0.06207 |
| L23 | 0.00053 | 0.47133 | 1.12536 | 0.29783 | 0.44675 |

**Table S6.** Brown–Forsythe tests comparing the variance of log-transformed centroid size between the first (F1) and tenth (F10) laboratory generations within each isofemale line. The table shows the variance in each generation, the F10/F1 variance ratio, the *F*-statistic, the unadjusted *P*-value, and the Benjamini–Hochberg-adjusted P-value. Ratios greater than 1 indicate greater variance in F10 than in F1.

| Line | $s^2$ F1 | $s^2$ F10 | F10/F1 | F | p_value | p_BH |
|---|---|---|---|---|---|---|
| L01 | 0.00111 | 0.00308 | 2.77100 | 4.53500 | 0.03770 | 0.05027 |
| L02 | 0.00073 | 0.00375 | 5.14800 | 18.02900 | 0.00008 | **0.00025** |
| L04 | 0.00042 | 0.00791 | 18.76800 | 10.52100 | 0.00235 | **0.00470** |
| L06 | 0.00028 | 0.00353 | 12.53900 | 4.99900 | 0.03037 | **0.04556** |
| L13 | 0.00103 | 0.00517 | 5.02500 | 2.79200 | 0.10100 | 0.12120 |
| L14 | 0.00063 | 0.00130 | 2.06000 | 1.28900 | 0.26090 | 0.28470 |
| L15 | 0.00338 | 0.00353 | 1.04500 | 0.16000 | 0.69110 | 0.69110 |
| L16 | 0.00040 | 0.00300 | 7.54500 | 11.54700 | 0.00123 | **0.00296** |
| L18 | 0.00030 | 0.00170 | 5.71600 | 21.01500 | 0.00003 | **0.00017** |
| L20 | 0.00047 | 0.00337 | 7.13400 | 20.34300 | 0.00004 | **0.00017** |
| L22 | 0.00024 | 0.00434 | 18.39100 | 44.69300 | 1.06E-08 | **1.27E-07** |
| L23 | 0.00053 | 0.00230 | 4.32300 | 9.75800 | 0.00327 | **0.00561** |

### Wing-shape analyses

**Table S7.** Procrustes linear-model comparisons of wing shape between the field sample (Field) and F1 of each isofemale line. A_PD is the adjusted Procrustes distance between the shapes predicted for Field and F1 at the joint mean log-transformed centroid size of the two groups. The table presents $R^2$, the F-statistic, effect size Z, and the Benjamini–Hochberg-adjusted P-value (p_BH) for the group effect. The columns prefixed with Ai describe the interaction between log-transformed centroid size and group, which tested whether the allometric trajectories differed between Field and F1; Ai_p_BH is the Benjamini–Hochberg-adjusted P-value for this interaction. In all comparisons, the joint mean log-transformed centroid size fell within the observed size ranges of both groups.

| Line | A_PD | $R^2$ | F | Z | p_BH | Ai_ $R^2$ | Ai_F | Ai_Z | Ai_p_BH |
|---|---|---|---|---|---|---|---|---|---|
| L01 | 0.0170 | 0.1170 | 5.8530 | 4.4610 | **0.0001** | 0.0240 | 1.2230 | 0.6280 | 0.7290 |
| L02 | 0.0220 | 0.2150 | 11.7830 | 5.4430 | **0.0001** | 0.0170 | 0.9360 | 0.0380 | 0.7290 |
| L04 | 0.0290 | 0.3000 | 12.7010 | 4.4010 | **0.0001** | 0.0280 | 1.1860 | 0.5530 | 0.7290 |
| L06 | 0.0260 | 0.2360 | 10.8230 | 4.5270 | **0.0001** | 0.0220 | 0.9950 | 0.1640 | 0.7290 |
| L13 | 0.0290 | 0.2440 | 14.2040 | 4.3800 | **0.0001** | 0.0140 | 0.7980 | -0.3040 | 0.8240 |
| L14 | 0.0190 | 0.1400 | 7.6320 | 4.3700 | **0.0001** | 0.0070 | 0.3610 | -1.8750 | 0.9710 |
| L15 | 0.0230 | 0.1920 | 7.1630 | 4.3900 | **0.0001** | 0.0270 | 0.9920 | 0.1450 | 0.7290 |
| L16 | 0.0130 | 0.0770 | 4.0080 | 3.0460 | **0.0014** | 0.0370 | 1.9250 | 1.5840 | 0.3380 |
| L18 | 0.0260 | 0.2890 | 17.1520 | 4.7030 | **0.0001** | 0.0470 | 2.7970 | 2.4420 | 0.0900 |
| L20 | 0.0200 | 0.2000 | 8.8910 | 4.9100 | **0.0001** | 0.0160 | 0.7030 | -0.5920 | 0.8660 |
| L22 | 0.0200 | 0.1380 | 7.4530 | 3.9920 | **0.0001** | 0.0090 | 0.4640 | -1.2280 | 0.9680 |
| L23 | 0.0380 | 0.3340 | 17.3170 | 4.2760 | **0.0001** | 0.0200 | 1.0550 | 0.3170 | 0.7290 |

**Table S8.** Procrustes linear-model comparisons of wing shape between F1 and F10 within each isofemale line. A_PD is the adjusted Procrustes distance between the shapes predicted for F1 and F10 at the selected reference value of log-transformed centroid size. The table presents $R^2$, the F-statistic, effect size Z, and the Benjamini–Hochberg-adjusted P-value (p_BH) for the group effect. The columns prefixed with Ai describe the interaction between log-transformed centroid size and group, which tested whether the allometric trajectories differed between F1 and F10; Ai_p and Ai_p_BH are the unadjusted and Benjamini–Hochberg-adjusted P-values for this interaction, respectively. Ref indicates whether the joint mean log-transformed centroid size fell within the observed ranges of both groups (T, true; F, false). When Ref = F but the size ranges overlapped (CS_O = Y), the midpoint of the shared range was used as the reference size. When the observed size ranges did not overlap (CS_O = N), the joint mean was retained and the comparison involved extrapolation. Such comparisons should be interpreted cautiously.

| Line | Ref | CS_O | A_PD | $R^2$ | F | Z | p_BH | Ai_ $R^2$ | Ai_F | Ai_Z | Ai_p | Ai_p_BH |
|---|---|---|---|---|---|---|---|---|---|---|---|---|
| L01 | T | Y | 0.021 | 0.143 | 10.287 | 5.120 | **0.0002** | 0.035 | 2.526 | 2.151 | 0.015 | **0.045** |
| L02 | F | Y | 0.019 | 0.086 | 5.669 | 3.740 | **0.0002** | 0.039 | 2.548 | 2.209 | 0.014 | **0.045** |
| L04 | T | Y | 0.022 | 0.177 | 9.578 | 5.207 | **0.0002** | 0.026 | 1.400 | 0.948 | 0.178 | 0.266 |
| L06 | F | Y | 0.018 | 0.049 | 2.923 | 2.763 | **0.0028** | 0.017 | 1.003 | 0.179 | 0.432 | 0.471 |
| L13 | T | Y | 0.017 | 0.107 | 6.253 | 4.287 | **0.0002** | 0.027 | 1.545 | 1.117 | 0.129 | 0.221 |
| L14 | T | Y | 0.018 | 0.056 | 3.802 | 2.967 | **0.0022** | 0.012 | 0.799 | -0.206 | 0.583 | 0.583 |
| L15 | T | Y | 0.032 | 0.083 | 5.824 | 4.302 | **0.0002** | 0.027 | 1.876 | 1.661 | 0.048 | 0.115 |
| L16 | T | Y | 0.026 | 0.069 | 4.881 | 3.403 | **0.0004** | 0.040 | 2.857 | 2.310 | 0.010 | **0.045** |
| L18 | F | N | 0.051 | 0.039 | 2.859 | 2.460 | **0.0075** | 0.041 | 2.946 | 2.497 | 0.006 | **0.045** |
| L20 | F | N | 0.031 | 0.132 | 10.609 | 4.243 | **0.0002** | 0.015 | 1.180 | 0.573 | 0.284 | 0.378 |
| L22 | T | Y | 0.034 | 0.231 | 21.500 | 4.365 | **0.0002** | 0.011 | 0.977 | 0.250 | 0.399 | 0.471 |
| L23 | F | Y | 0.019 | 0.107 | 5.208 | 3.920 | **0.0002** | 0.034 | 1.648 | 1.241 | 0.111 | 0.221 |

**Table S9.** Observed total Procrustes variance and bootstrap estimates shown in Figure 4A for the field sample of males (Field) and males from the first (F1) and tenth (F10) generations maintained in the laboratory. Bootstrap estimates were based on 9,999 replicates, each containing 17 individuals sampled with replacement from the corresponding group. Observed variance (Observed $s^2$) was calculated using all available individuals. Lower 95% and Upper 95% indicate the 2.5th and 97.5th percentiles of the bootstrap distribution. Analyses were performed on Procrustes coordinates without allometric correction.

| Group | N | Observed $s^2$ | Bootstrap_median | Lower_95 | Upper_95 |
|---|---|---|---|---|---|
| Field | 17 | 4.14E-04 | 3.90E-04 | 3.08E-04 | 4.70E-04 |
| F1 | 267 | 5.77E-04 | 5.69E-04 | 4.28E-04 | 7.51E-04 |
| F10 | 360 | 8.18E-04 | 8.03E-04 | 5.83E-04 | 1.11E-03 |

**Table S10.** Pairwise bootstrap comparisons of total Procrustes variance among the field sample of males (Field) and males from the first (F1) and tenth (F10) generations maintained in the laboratory. Differences were calculated as the variance of the first group listed minus that of the second, whereas ratios were calculated by dividing the variance of the first group by that of the second. Med. diff. and Med. ratio indicate the medians of the bootstrap distributions. Lower_95 and Upper_95 indicate the lower and upper limits of the 95% bootstrap interval, respectively. Differences were considered supported when the interval did not include zero, and variance ratios were considered supported when the interval did not include one. Estimates were based on 9,999 bootstrap replicates, each containing 17 individuals sampled with replacement from each group. Analyses were performed on Procrustes coordinates without allometric correction.

| Comp. | Med. diff. | Lower_95 | Upper_95 | Med. ratio | Lower_95 | Upper_95 |
|---|---|---|---|---|---|---|
| F1 – Field | 1.795E-04 | 1.681E-05 | 3.779E-04 | 1.4624 | 1.0395 | 2.1031 |
| F10 – Field | 4.157E-04 | 1.757E-04 | 7.256E-04 | 2.0673 | 1.4171 | 3.0683 |
| F10 – F1 | 2.336E-04 | -5.414E-05 | 5.716E-04 | 1.4161 | 0.9229 | 2.1870 |

**Table S11**. Bootstrap comparisons of within-line Procrustes variance among the field sample of males (Field) and males from the first (F1) and tenth (F10) laboratory generations of each isofemale line. Comparisons were calculated as the variance of the first group listed minus that of the second. Med. $s^2$ A and Med. $s^2$ B indicate the median bootstrap variances of the first and second groups, respectively, and Med. diff. is the median bootstrap difference. Lower_95 and Upper_95 indicate the 2.5th and 97.5th percentiles of the bootstrap distribution of the differences. Estimates were based on 9,999 bootstrap replicates, each containing 17 individuals sampled with replacement from each group. Two-sided empirical bootstrap P-values were calculated from the proportions of bootstrap variance differences on either side of zero and adjusted using the Benjamini–Hochberg procedure separately within each set of 12 comparisons. The 95% bootstrap intervals were not adjusted for multiple comparisons.

| Comparison | Line | $N_A$ | $N_B$ | Med $s^2_A$ | Med. $S^2_B$ | Med. Diff | Lower_95 | Upper_95 | pBH |
|---|---|---|---|---|---|---|---|---|---|
| F1 – Field | L01 | 27 | 17 | 4.95E-04 | 3.89E-04 | 1.06E-04 | -3.78E-05 | 2.72E-04 | 0.54 |
| | L02 | 28 | 17 | 4.02E-04 | 3.90E-04 | 1.23E-05 | -1.21E-04 | 1.51E-04 | 0.94 |
| | L04 | 13 | 17 | 3.55E-04 | 3.90E-04 | -3.54E-05 | -1.42E-04 | 7.04E-05 | 0.87 |
| | L06 | 17 | 17 | 3.84E-04 | 3.89E-04 | -4.85E-06 | -1.25E-04 | 1.18E-04 | 0.94 |
| | L13 | 22 | 17 | 4.80E-04 | 3.90E-04 | 9.14E-05 | -3.95E-05 | 2.40E-04 | 0.54 |
| | L14 | 30 | 17 | 4.15E-04 | 3.90E-04 | 2.58E-05 | -1.03E-04 | 1.59E-04 | 0.94 |
| | L15 | 13 | 17 | 4.98E-04 | 3.89E-04 | 1.09E-04 | -3.98E-05 | 2.48E-04 | 0.54 |
| | L16 | 30 | 17 | 4.56E-04 | 3.91E-04 | 6.59E-05 | -6.28E-05 | 2.06E-04 | 0.64 |
| | L18 | 25 | 17 | 3.82E-04 | 3.90E-04 | -5.65E-06 | -1.36E-04 | 1.56E-04 | 0.94 |
| | L20 | 20 | 17 | 3.04E-04 | 3.90E-04 | -8.56E-05 | -2.00E-04 | 3.09E-05 | 0.54 |
| | L22 | 29 | 17 | 4.91E-04 | 3.90E-04 | 1.03E-04 | -7.27E-05 | 2.82E-04 | 0.63 |
| | L23 | 13 | 17 | 3.67E-04 | 3.90E-04 | -2.28E-05 | -1.51E-04 | 1.05E-04 | 0.94 |
| F10 – Field | L01 | 30 | 17 | 4.02E-04 | 3.90E-04 | 1.33E-05 | -1.03E-04 | 1.38E-04 | 0.89 |
| | L02 | 30 | 17 | 4.71E-04 | 3.90E-04 | 8.18E-05 | -4.21E-05 | 2.18E-04 | 0.82 |
| | L04 | 30 | 17 | 5.20E-04 | 3.90E-04 | 1.30E-04 | -2.14E-05 | 2.87E-04 | 0.56 |
| | L06 | 30 | 17 | 4.41E-04 | 3.90E-04 | 5.29E-05 | -9.03E-05 | 2.20E-04 | 0.82 |
| | L13 | 30 | 17 | 4.49E-04 | 3.90E-04 | 6.03E-05 | -9.53E-05 | 2.54E-04 | 0.82 |
| | L14 | 30 | 17 | 4.35E-04 | 3.89E-04 | 4.77E-05 | -1.14E-04 | 2.68E-04 | 0.82 |
| | L15 | 30 | 17 | 4.44E-04 | 3.90E-04 | 5.39E-05 | -8.61E-05 | 2.00E-04 | 0.82 |
| | L16 | 30 | 17 | 4.09E-04 | 3.89E-04 | 2.06E-05 | -1.06E-04 | 1.64E-04 | 0.89 |
| | L18 | 30 | 17 | 3.54E-04 | 3.90E-04 | -3.44E-05 | -1.51E-04 | 8.86E-05 | 0.82 |
| | L20 | 30 | 17 | 3.30E-04 | 3.90E-04 | -5.82E-05 | -1.68E-04 | 6.45E-05 | 0.82 |
| | L22 | 30 | 17 | 5.74E-04 | 3.90E-04 | 1.84E-04 | 1.50E-05 | 3.68E-04 | 0.4 |
| | L23 | 30 | 17 | 4.01E-04 | 3.90E-04 | 1.38E-05 | -1.37E-04 | 2.21E-04 | 0.89 |
| F10 – F1 | L01 | 30 | 27 | 4.02E-04 | 4.95E-04 | -9.41E-05 | -2.56E-04 | 5.76E-05 | 0.82 |
| | L02 | 30 | 28 | 4.70E-04 | 4.02E-04 | 6.77E-05 | -7.99E-05 | 2.13E-04 | 0.82 |
| | L04 | 30 | 13 | 5.19E-04 | 3.55E-04 | 1.65E-04 | 2.01E-05 | 3.17E-04 | 0.31 |
| | L06 | 30 | 17 | 4.43E-04 | 3.85E-04 | 5.82E-05 | -8.99E-05 | 2.25E-04 | 0.82 |
| | L13 | 30 | 22 | 4.46E-04 | 4.79E-04 | -3.04E-05 | -2.17E-04 | 1.80E-04 | 0.82 |
| | L14 | 30 | 30 | 4.36E-04 | 4.15E-04 | 2.35E-05 | -1.53E-04 | 2.46E-04 | 0.82 |
| | L15 | 30 | 13 | 4.45E-04 | 4.98E-04 | -5.35E-05 | -2.19E-04 | 1.18E-04 | 0.82 |
| | L16 | 30 | 30 | 4.10E-04 | 4.53E-04 | -4.40E-05 | -1.99E-04 | 1.08E-04 | 0.82 |
| | L18 | 30 | 25 | 3.53E-04 | 3.81E-04 | -2.83E-05 | -1.95E-04 | 1.12E-04 | 0.82 |
| | L20 | 30 | 20 | 3.30E-04 | 3.03E-04 | 2.89E-05 | -8.47E-05 | 1.47E-04 | 0.82 |
| | L22 | 30 | 29 | 5.76E-04 | 4.95E-04 | 8.13E-05 | -1.35E-04 | 3.06E-04 | 0.82 |
| | L23 | 30 | 13 | 4.01E-04 | 3.68E-04 | 3.60E-05 | -1.24E-04 | 2.48E-04 | 0.82 |

**Table S12.** Observed total Procrustes variance for Field, F1, and F10, and its decomposition into within-line and between-line components for the two laboratory generations. Total $s^2$ indicates the observed total Procrustes variance, whereas w $s^2$ and b $s^2$ indicate the within-line and between-line components, respectively. Prop. w and Prop. b represent the proportions of total variance attributable to the within-line and between-line components. The field sample (Field) was not decomposed because it did not consist of established laboratory lines. Analyses were performed on Procrustes coordinates without allometric correction.

| Group | Total $s^2$ | w $s^2$ | b $s^2$ | Prop. w | Prop. b |
|---|---|---|---|---|---|
| Field | 0.00041 | - | - | - | - |
| F1 | 0.00058 | 0.00043 | 0.00015 | 0.74313 | 0.25687 |
| F10 | 0.00082 | 0.00044 | 0.00038 | 0.53903 | 0.46097 |

**Table S13.** Sensitivity analysis of Procrustes variance components to differences in sample size between F1 and F10. In each of 9,999 subsampling replicates, 13 individuals were randomly selected without replacement from each of the 12 isofemale lines in both generations (N = 156 per generation). Differences were calculated as F10 minus F1. Median represents the median difference, P2.5 and P97.5 represent the corresponding percentiles of the subsampling distributions, and Proportion positive indicates the proportion of replicates in which the variance component was greater in F10 than in F1. Analyses were performed on Procrustes coordinates without allometric correction. The median between-line variance ratio (F10/F1) was 2.28 (2.5th–97.5th percentiles: 1.95–2.66).

| Component | Median difference | P2.5 | P97.5 | Proportion positive |
|---|---|---|---|---|
| Total | $2.34429 \times 10^{-4}$ | $1.69841 \times 10^{-4}$ | $2.99087 \times 10^{-4}$ | 1 |
| Within-line | $1.10648 \times 10^{-5}$ | $-2.82598 \times 10^{-5}$ | $5.19636 \times 10^{-5}$ | 0.713 |
| Between-line | $2.22812 \times 10^{-4}$ | $1.75033 \times 10^{-4}$ | $2.74294 \times 10^{-4}$ | 1 |
